# Pericentromeric heterochromatin impacts genome compartmentalization and sex chromosome evolution in a fish

**DOI:** 10.1101/2021.03.01.433482

**Authors:** Lingzhan Xue, Yu Gao, Meiying Wu, Haiping Fan, Yongji Huang, Zhen Huang, Dapeng Li, Luohao Xu

## Abstract

Compartmentalization is one of the principles of chromosome 3D organization and has been suggested to be driven by the attraction of heterochromatin. The extent to which the pericentromeric heterochromatin (PCH) impacts chromosome compartmentalization is yet unclear. Here we produced a chromosome-level and fully phased diploid genome of an aquaculture fish, zig-zag eel (*Mastacembelus armatus*), and identified the centromeric and pericentromeric regions in the majority of chromosomes of both haploid genomes. The PCH is on average 4.2 Mb long, covering 17.7% of the chromosomes, and is the major target of histone 3 lysine 9 trimethylation (H3K9me3). In nearly half of the chromosomes, the PCH drives the chromosomes into two or three megascale chromatin domains with the PCH being a single one. We further demonstrate that PCH has a major impact in submetacentric, metacentric and small telocentric chromosomes in which the PCH drives the distribution of active and inactive compartments along the chromosomes. Additionally, we identified the young and homomorphic XY sex chromosomes that are submetacentric with the entire short-arm heterochromatinized. Interestingly, the sex-determining region seems to arise within the PCH that has been in place prior to the X-Y divergence and recombination suppression. Together, we demonstrate that the PCH can cover a considerably large portion of the chromosomes, and when it does so, it drives chromosome compartmentalization; and we propose a new model for the origin and evolution of homomorphic sex chromosomes in fish.

## Introduction

The eukaryotic chromosomes are highly folded and organized into three-dimensional space, with different parts of the chromosomes exposed to different nuclear environments (Bonev and Cavalli 2016; van Steensel and Belmont 2017). The chromatin is typically divided into active (A) and inactive (B) compartments, and the same class of compartments tends to interact with each other (Lieberman-Aiden et al. 2009). Recent studies have highlighted the role of liquid-liquid phase separation in driving the organization of compartments, primarily due to attraction of multivalent molecules including HP1 protein, H3K9me3 and H3K27me3, among others (Larson et al. 2017; Strom et al. 2017). In particular, a recent study on inverted nuclei suggests that attraction of heterochromatin is the major factor determining the genome compartmentalization (Falk et al. 2019).

Centromeres and the pericentromeric regions have often been thought as a sink of constitutive heterochromatin, harboring the major H3K9me3 targets (Pidoux and Allshire 2005; Saksouk, Simboeck, and Déjardin 2015; Lehnertz et al. 2003). Moreover, centromeres and pericentromeric heterochromatin (PCH) are often localised near nuclear lamina where the chromatin are largely silenced (Guelen et al. 2008; van Steensel and Belmont 2017), and in some taxa they may cluster together to form chromocenter (Jagannathan, Cummings, and Yamashita 2018; Fransz et al. 2002; Jones 1970). Unfortunately, the centromeres are composed of large arrays of tandem repeats and the pericentromeric regions are rich in transposable elements and satellite repeats (Hartley and O’Neill 2019; Sullivan, Chew, and Sullivan 2017; Melters et al. 2013), and their repetitive nature have impeded the study on their functions and structures until recently (Miga et al. 2020; Jain et al. 2018; Miga 2020). Long-read sequencing technology, particularly ultralong long Nanopore sequencing, has recently demonstrated its power in assembling the complete sequence of human centromeres (Bzikadze and Pevzner 2020). The low accuracy of Nanopore reads, however, required careful polishing and manual curation for the assembled centromere sequences.

In this study, we took advantage of the HiFi sequencing technology (Wenger et al. 2019) to produce long and accurate reads of an aquaculture fish zig-zag eel (*Mastacembelus armatus*). The fully phased and chromosome levels assembly of the diploid genomes are highly continuous, including centromeric satellite sequences for most chromosomes in the two haploid genomes. The almost complete genomes provide us an unprecedented opportunity to study the role of centromeric and pericentromeric heterochromatin in genome compartmentalization.

## Result

### Fully phased chromosome-level assembly

The CCS reads have a long read length (~15 kb) and our *k-mer* analysis suggests the error rate is only 0.086% (**Fig. 1a**, **Supplementary Fig. S1**). The long and accurate reads allow for a more accurate estimate of the genome size (600.1 Mb) because of the use of a larger k-*mer* size (256 in this case), and are useful to assemble the fully phased diploid genomes. The principle of assembling two haploid genomes is to first produce a collapsed (or pseudo-haploid) genome assembly which will be mapped by the CCS reads, followed by the partitioning of haplotype-specific reads for separate genome assembly (**Fig. 1b**). The draft collapsed assembly is already highly continuous, with only 228 contigs and a contig N50 value of 9.9 Mb. We then linked the haploid contigs into chromosomes according to their anchoring positions in the existing chromosomal assembly of zig-zag eel (fMasArm1.2) (Rhie et al. 2020). The CCS reads were then mapped against the pseudo-chromosomal genome and were phased by combining the phasing information derived from CCS reads and Hi-C read pairs sequenced from the same individual (**Fig. 2b**). On average the largest phased blocks span 99.6% of the pseudo-chromosomes, suggesting the chromosomes were able to be nearly completely phased for the two haplotype (**Supplementary Table 1**).

**Fig. 1.**
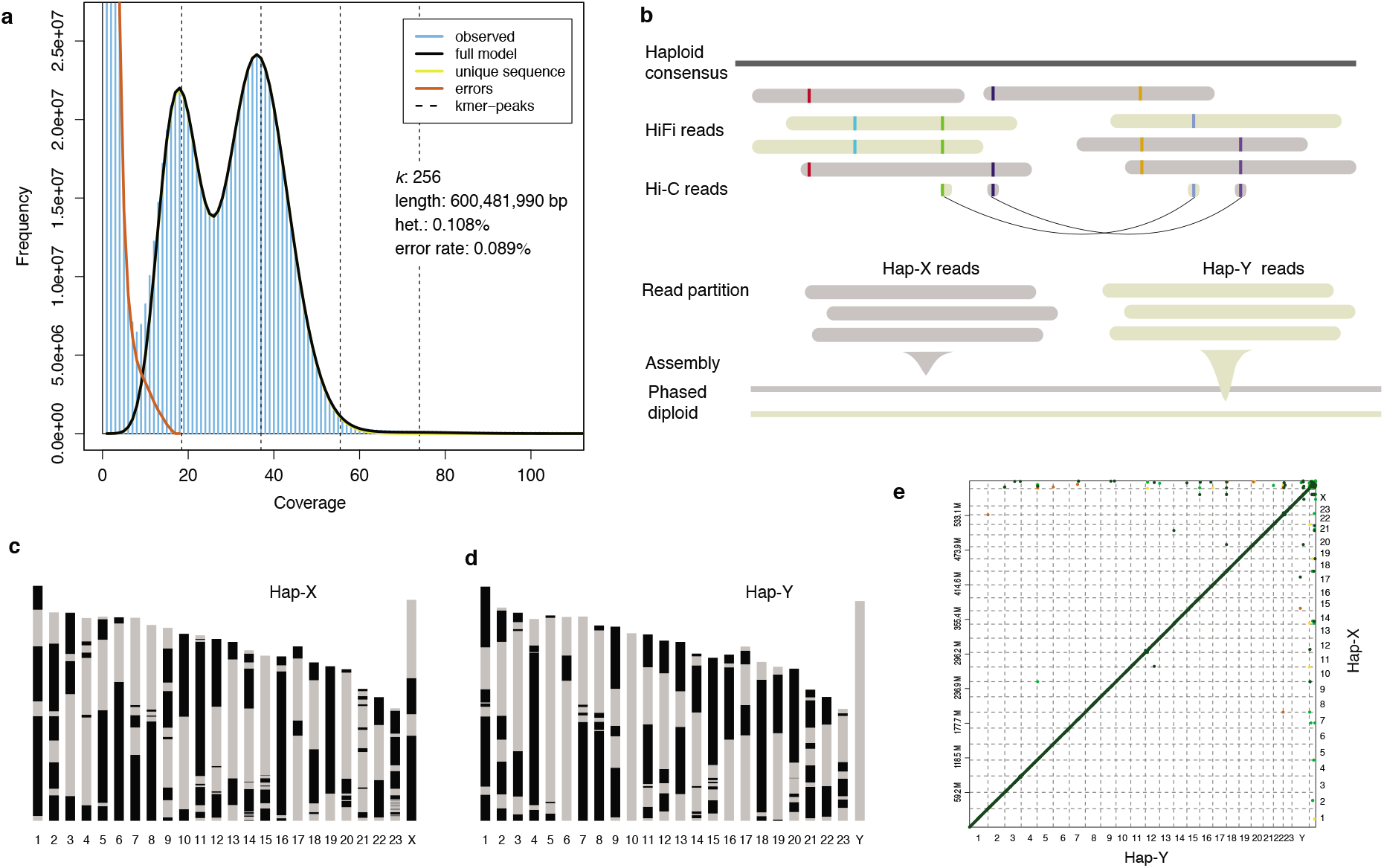
Phased diploid genome assembly. **a)** Estimation of genome size and heterozygosity by the distribution of *k*-mers (k = 256) of HiFi reads. The haploid peak at ~19X is nearly as high as the dipoid peak despite. **b)** The workflow of phased diploid genome assembly. The haploid consensus was generated by the Peregrine assembler. Both the HiFi reads and Hi-C reads were used to phase and partition the haplotype-specific HiFi reads. The two haplotypes are named as Hap-X and Hap-Y. **c-d)** The assembled chromosomes of both Hap-X and Hap-Y genomes. One color (black or grey) represents a continuous sequence (contig). The Y chromosome has no gap. **e)** The genomic synteny between the Hap-X and Hap-Y genome assemblies. The unanchored scaffolds are shown at the right and top edges.

**Fig. 2.**
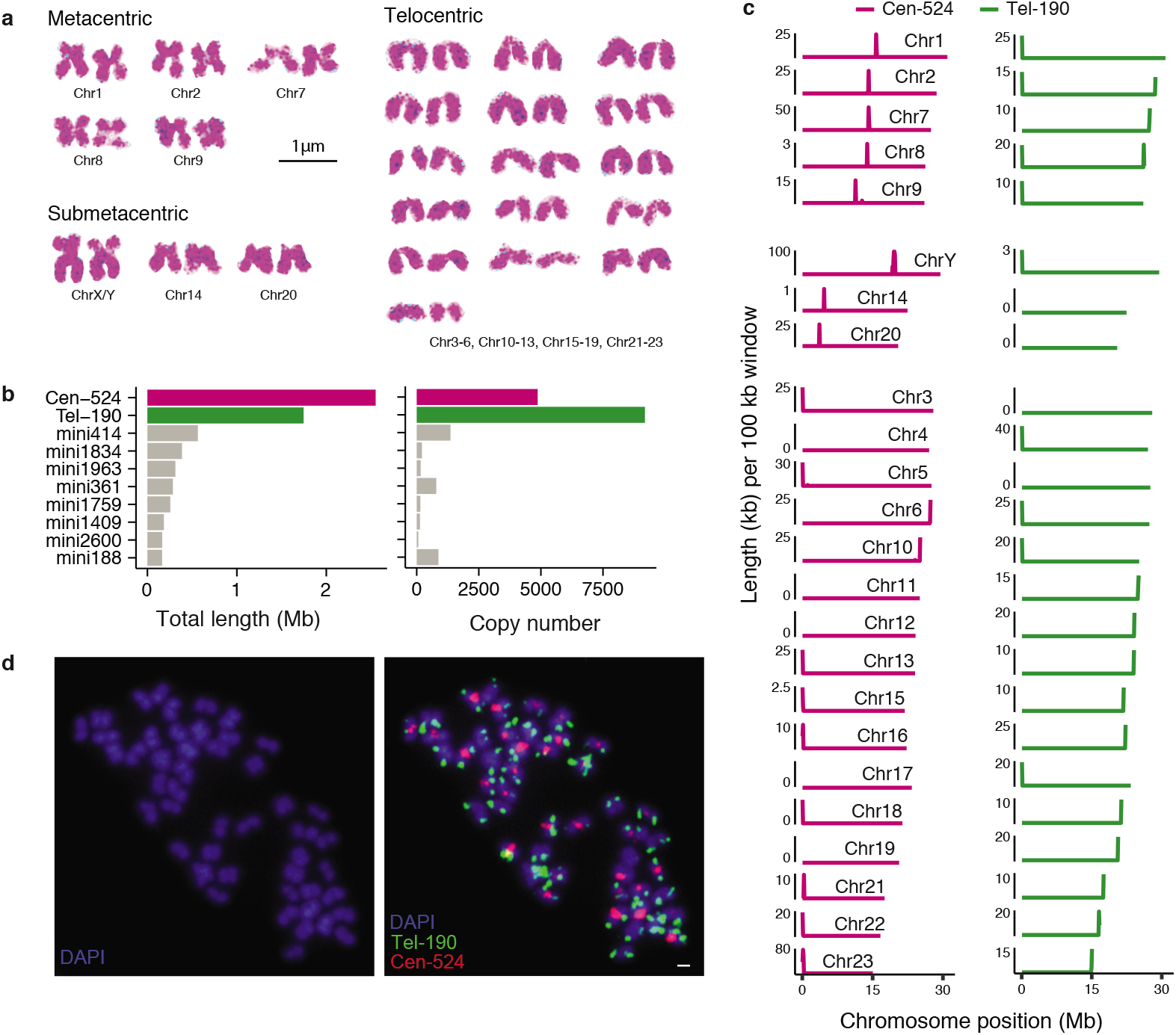
The karyotype and the identification of centromeres. **a)** The chromosomes were grouped according to the chromosome types. The chromosome IDs were assigned according the size of the assembled chromosomes. Most telocentric chromosome cannot be differentiated. **b)** The total length and copy number of the top 10 abundant satellite DNA. The putative centromeric repeats (Cen-524 and Tel-190) were highlighted in color. **c)** The distribution of Cen-524 and Tel-190 satellite DNA along the chromosomes. The chromosomes were grouped according to their morphological classification in a). In a few chromosomes (e.g. Chr4) the satellite DNA sequence have not been assembled. **d)** The FISH results of Cen-524 (red) and Tel-190 (green) on the chromosomes. The scale of the white bar is 1μm.

The partitioned haplotype-specific CCS reads were assembled into two haploid genomes, named as hap-X and hap-Y respectively (**Fig. 2b**). Both haploid genomes have comparable genome size and contig N50 values with the haploid consensus assembly (**Table 1**). We further extracted the hap-X and hap-Y specific Hi-C read-pairs, and used them to link the hap-X and hap-Y contigs into chromosomes, respectively. Hence, we produced two independently assembled chromosome-level haploid genomes (**Fig. 2c-d**) between which we did not detect visible large-scale chromosome rearrangements (**Fig. 2e**), but detected a few inversions near telomeres between our new assembly and the previous assembly (**Supplementary Fig. S2**). In the following analyses we use the hap-Y assembly as the representative haploid genome unless stated otherwise.

**Table 1.** The haploid and diploid genome assembly

| Assembly |  | Size (Mb) | Contig N50 (Mb) | # contig | BUSCO (%) |
| --- | --- | --- | --- | --- | --- |
| Haploid consensus |  | 595.7 | 9.9 | 474 | 94.6 |
| Diploid | Hap-X | 582 | 7.8 | 364 | 94.4 |
|  | Hap-Y | 585.6 | 8.6 | 366 | 97.1 |

### Phylogenomics

Using the whole genome data of seven Percomorpha species and one outgroup species *Acanthochaenus luetkenii* (a basal Acanthopterygii fish) (Musilova et al. 2019), we placed zig-zag eel as a close-related species of the Asian swamp eel (*Monopterus albus*), in the same order Synbranchiformes (Tian, Hu, and Li 2021). We further dated the divergence among those fish species, and estimated zig-zag eel and Asian swamp eel diverged from each other ~36 million years ago (**Supplementary Fig. S3**).

### Percomorpha karyotype

The karyotype remains relatively stable in Percomorpha with most species having a chromosome number of 48 (2n), but the variation of chromosome number has been observed in many taxa (Paim et al. 2017; Motta-Neto et al. 2019). We took advantage of two additional chromosome-level genome assemblies of Percomorpha fish with varying chromosome number: Nile tilapia (2n = 44) (Tao et al. 2021) and big-bellied seahorse (2n = 42), to reconstruct the Percomorpha ancestral karyotype. Using a parsimonious approach, we inferred three fusions having occurred in big-belly seahorse and two fusions in Nile tilapia (**Supplementary Fig. S4**). Those inferred fusion events are sufficient to explain the difference in chromosome number among the three Percomorpha species, confirming haploid number of 24 is likely an ancestral feature of Percomorpha. Like other Percomorpha species, zig-zag eel does not possess microchromosomes, with the smallest chromosome (15.2 Mb) being only half-size of the largest chromosome (31.4 Mb) (**Fig. 1c-d**). There are five metacentric chromosomes that are all among the top nine largest chromosomes; three chromosome pairs are submetacentric and the rest 16 chromosome pairs are telocentric (**Fig. 2a**).

### Genomic and cytogenetic identification of centromeric satellites

To identify the centromeric satellite DNA, we set out to look for the most abundant satellite sequences in the assembled genome (Melters et al. 2013). Two satellite sequences show prominent sequence amount and copy number compared to the others, having a monomer length of 524 and 190 bp respectively (**Fig. 2b**). Interestingly, the 524-bp satellite (Cen-524) usually appears at one single loci on a chromosome, and in telocentric chromosome it appears at the chromosomal ends while in metacentric and submetacentric it appears in the middle (**Fig. 2c**). This makes Cen-524 a strong candidate for the centromeric satellite. The 190-bp satellite (Tel-190), on the other hand, appears exclusively at the chromosomal ends, and on metacentric chromosomes it sometimes appears at both ends of the chromosomes (**Fig. 2c**), suggesting Tel-190 is associated with telomeres rather than centromeres. To further validate the candidate centromeric satellite, we hybridized the probes of Cen-524 and Tel-190 using the fluorescent in situ hybridization (FISH) technique on the chromosomes, and found their locations on the chromosomes (**Fig. 2d**) are largely consistent with the genomic sequence assembly. We have unfortunately not been able to assemble the conserved telomeric motif (TTAGGG)n (Meyne, Ratliff, and Moyzis 1989) for most chromosomes, likely because the ends of contigs at chromosomal tips break with long arrays of Tel-190 sequences, but our FISH experiments nevertheless show the co-presence of the (TTAGGG)s motifs and Tel-190 (**Supplementary Fig. S5**).

### Pericentromeric heterochromatin

The average size of the assembled core centromeres (satellite DNA) is ~27.1 kb, but the pericentromeric regions rich in repetitive sequences appear to be much larger. To demarcate the boundaries of pericentromeric regions, we examined the landscape of repeat abundance on the chromosomes. It is readily recognized that a considerably large region (~4 Mb) around the centromeres has an elevated repeat content of typically higher than 50%, compared to that of the other part of the genomes (less than 16%) (**Fig. 3a-c, Supplementary Table S2**). Furthermore, the highly-repetitive regions have a lower gene density, lower recombination rate and more frequent H3k9me3 modifications (**Fig. 3a-c**, **Supplementary Fig. S6-9**), making them the putative pericentromeric regions. Combining the information of repetitive levels, gene density, recombination rate and H3K9me3 modification, we carefully demarcate the boundary of the pericentromeric region in each chromosome. Because the entire pericentromeric regions show a high degree of H3K9me3 modification and accumulation of repetitive sequences, they were further considered as pericentromeric heterochromatin (PCH). The total PCH accounts for 53.7% of the genome-wide H3K9me3 peaks (**Supplementary Fig. S6-9**), though it covers only 17.7% of the genome.

**Fig. 3.**
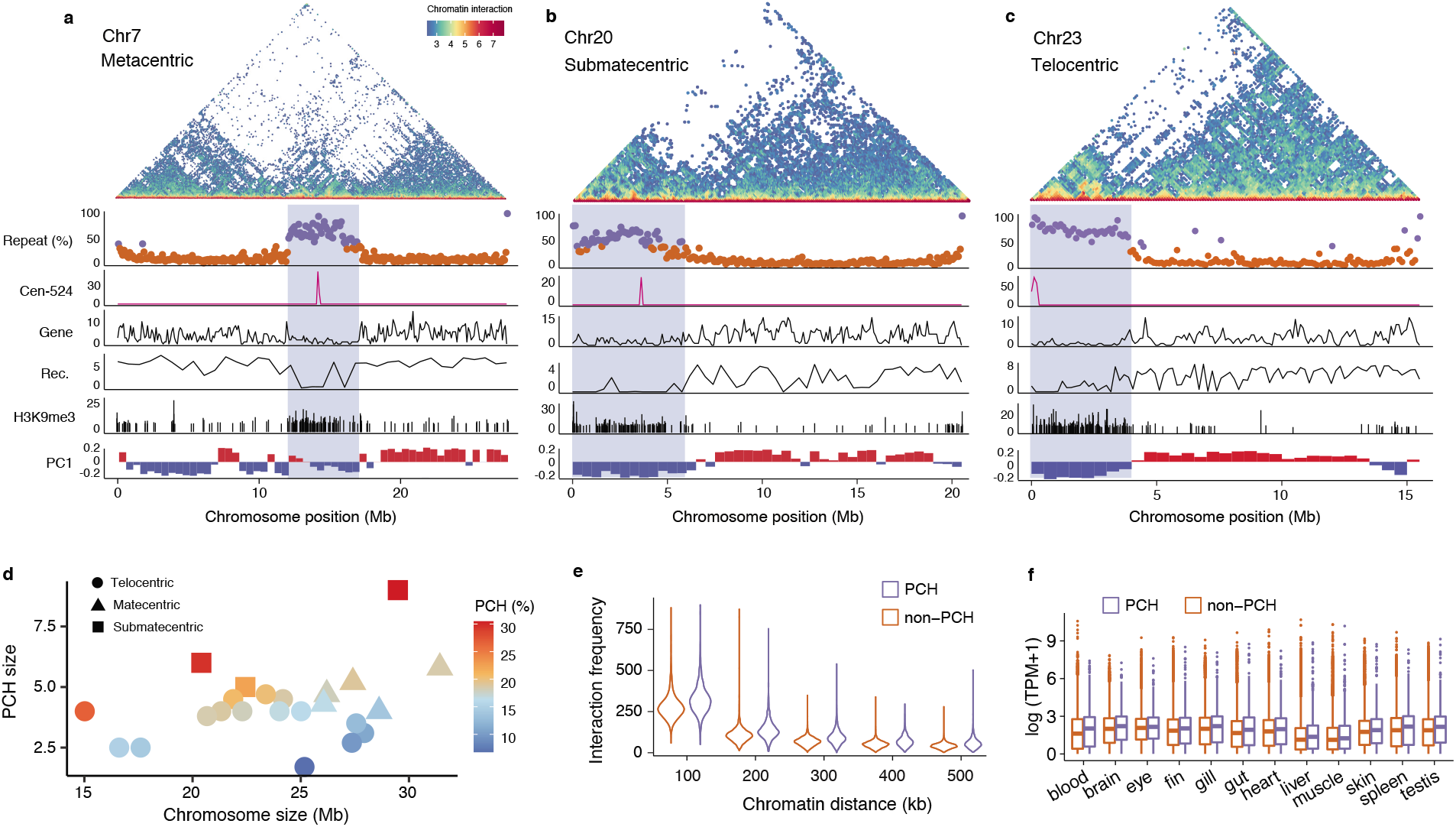
The impact of PCH on genome compartmentalization. **a-c)** Three chromosomes (Chr7, Chr20, Chr23) representing the metacentric, submetacentric and telocentric chromosomes, respectively. In the top panel, the colors of dots measure the frequency of chromatin interacting between 100 kb windows. When the repeat content of a 50 kb sequence (a dot) is larger than 40%, it is highlighted in dark purple, otherwise in orange. The portion (%) of Cen-524 satellite DNA in 100 kb windows is shown in pink. The peaks represent the locations of centromeres. The gene density is measured as the number of genes in 100 kb windows. The recombination rate (Rec.) is estimated with selected window size based on the available variants. The Y-axis of the H3K9me3 panel shows the −log 10 transformed p-values for the H3K9me3 peaks. The PC1 panel shows the PC1 values of Hi-C epivector: the positive values (red) represent active (A) compartments and the negative values (blue) represent inactive (B) compartments. The PCH regions are highlighted with blue background **d)** Weak positive correlation of chromosome size and PCH size. The colors measures the proportions of PCH on the chromosomes. **e)** Interaction is more frequent between PCH chromatin over various genomic distance than between non-PCH. **f)** Gene are expressed at a higher level in PCH than in non-PCH.

The sizes of PCH show only weak positive correlation with chromosome sizes without statistical significance (Pearson’s r = 0.34, P = 0.11), and the majority of PCH has a size around 4.2 Mb (**Fig. 3d**). As a consequence, smaller chromosomes, particularly for the telocentric ones, tend to possess a larger portion of PCH (**Fig. 3d**). Because almost the entire short-arm of submetacentric chromosomes have been heterochromatinized (**Supplementary Fig. S6**), this type of chromosomes have larger PCH than metacentric or telocentric chromosomes (**Fig. 3d**). Within PCH, chromatin interaction is more frequent than within non-PCH over various large genomic distances, consistent with their higher degree of folding and compaction (**Fig. 3e**). Unexpectedly, PCH regions have a larger portion of active genes whose expression has a high level and breadth than those in non-PCH (**Fig. 3f**, **Supplementary Fig. S10**). This implies the partial role of H3K9me3 in gene repression and the likely presence of other epigenetic modifications that activate the expression of the genes within the PCH (Feng et al. 2020; Burton et al. 2020; Saksouk, Simboeck, and Déjardin 2015).

### Impact of PCH on chromosome compartmentalization

One characteristic of PCH is a wide spread of H3K9me3 histone modification that is expected to drive the chromatin into distinct compartments from the other part of the genome (Bian et al. 2020; L. Wang et al. 2019). When the proportion of the PCH on the chromosome reach as high as 25%, the impact of PCH on chromatin compartmentalization can extent to the entire chromosome, but the extend of the impact varies among chromosome depending on the chromosome size and the position of centromere. For metacentric chromosomes, the chromosomes are typically divided into three megascale domains, with the PCH forming a single small domain at the center and the two chromosome arms forming two domains (**Fig. 3a**, **Supplementary Fig. S6**). This is consistent with previous understanding that the centromere often serves as the chromatin barrier between the two arms (Muller, Gil, and Drinnenberg 2019). Interestingly, it’s frequently observed that one of the arms tend to be embedded into active compartments while the other tend to form inactive compartments, except for the largest chromosome (Chr1) (**Fig. 3a**, **Supplementary Fig. S6**).

For submetacentric chromosomes, the chromosomes are roughly divided into two domains, with the PCH forming a single inactive compartment while the rest part of the chromosome tending to form active compartments (**Fig. 3b**, **Supplementary Fig. S7**). When the chromosomes are large (e.g chrY), the PCH spreads only within the short arm, but in the smaller chromosome (chr20), the PCH can extend beyond the centromere into the long arm (**Fig. 3b**, **Supplementary Fig. S7**).

For telocentric chromosomes, the PCH is usually smaller; but in smaller chromosomes when the proportions of PCH become larger, the impact of PCH imposing on chromosome compartmentalization becomes more significant. Specifically, in chr15, chr17, chr22 and chr23, the average proportion of PCH is 20.1% which drives the chromosomes into two large domains (**Fig. 3c**, **Supplementary Fig. 8**). Additionally, the chromosome compartmentalization results in an B-A-B structure, with the PCH and the other chromosome tip forming single B compartments and the rest chromosome body being an A compartment (**Fig. 3c**, **Supplementary Fig. 8**). In the other telocentric chromosomes where the average proportion of PCH is 14.6%, frequent transitions between A and B compartments along the chromosomes are observed (**Supplementary Fig. 9**).

### Young sex chromosome

Zig-zag eel is a sequentially hermaphroditic fish in which all individuals initially develop as females but about half of them later develop into males. A recent study, interestingly, identified male-specific markers, implying the existence of a sex chromosome pair (XY) in the genome (Xue et al. 2020). To identify the sex chromosomes, we collected re-sequencing data from 10 males and 10 females, and screened for the sex-linked markers. A ~7 Mb sequence in the second largest chromosome was found to be linked with sex (**Fig. 4a**), displaying male-specific variants (**Fig. 4b-c**) and large population divergence between males and females (**Supplementary Fig. S11**). The chromosome containing male-specific variants was therefore named as the Y chromosome (in the hap-Y assembly) whereas the homozygous chromosome was named as the X chromosome (in the hap-X assembly). The sex-determining (SD) region spans the centromere, but the majority of its sequence is within the PCH (**Fig. 4c-d**). Despite the large size of the SD region, the divergence of the X and Y is extremely low, with the X and Y sequence similarity remaining at 99% (**Fig. 4d**). Moreover, we did not detect differences in gene content between the X and Y chromosome in the SD region.

**Fig. 4.**
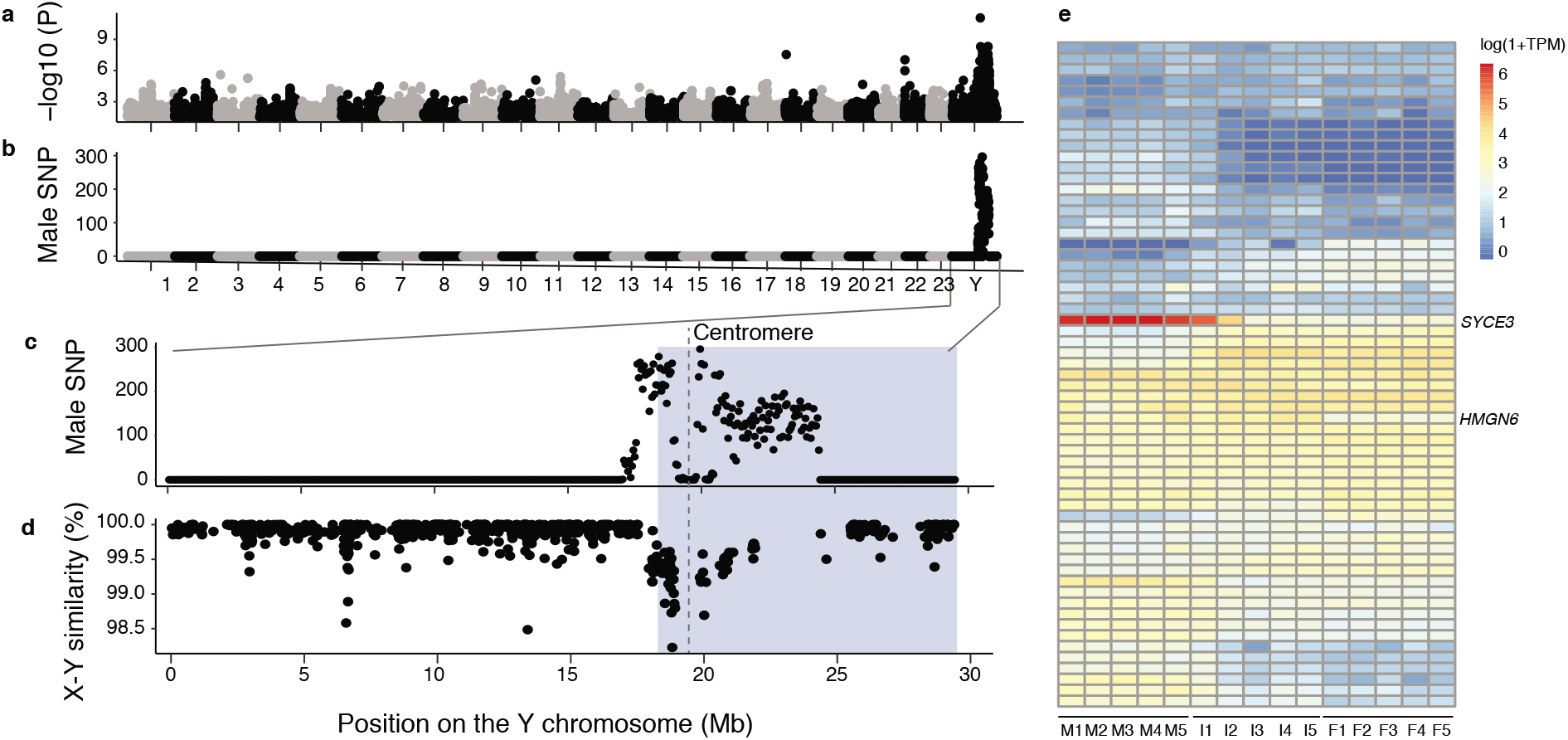
Identification of the sex-linked region and genes. **a)** p-values (log10 transformed) for genome-wide association study for the sex trait (male or female). Re-sequencing data of ten male and ten female individuals were used. **b)** The density (number per 50 kb windows) of male-specific SNPs. Those SNPs are present in all ten males but not in females. **c)** The zoom-in view of b) on the Y chromosome. The vertical dashed line denote the position of centromere. PCH is highlighted with blue background. **d)** The sequence similarity between the introns of the X and Y chromosomes. The similarity was calculated in every 100 kb window (black dot). Gene deserts were frequently seen in the sex-linked region. **e)** expression profiling of the gonads of five individuals of male (M), female (F) and intersex (I). *SYCE3* has a testis-specific expression and *HMGN6* is expressed in both testis and ovotestis.

To identify genes that are responsible for sex reversal, we collected gonadal transcriptomes from males and females as well as the intersex individuals. Among the 92 genes in the SD region, 59 are expressed. In particular, gene *SYCE3* has a significantly high expression in testis and the late-stage of ovotestis (I2 and I4) (**Fig. 4e**). In addition, *HMGN6* is expressed in testis and every stage of ovotestis at a similar level, but at much lower level in ovary (**Fig. 4e**). This suggests the role of *HMGN6* in development and maintenance of testis but *SYCE3* may be involved in spermatogenesis or other biological processes in mature testis.

### New model of sex chromosome origin

A lack of inversions between the non-recombining regions of X and Y chromosome implies that the suppression of recombination is not mechanistic (**Fig. 5a**). Though the non-recombining region is as large as ~7 Mb, it’s mainly in the PCH, spanning around the centromere. We further found that similar to the Y chromosome, the X chromosome has a large PCH occupying almost the entire short arm (**Fig. 5b-c**). This suggests the extensive spreading to PCH across the short arm of the Y chromosome is not the consequence of recombination arrest, rather, the PCH has already fully expanded in the proto-sex chromosome. If this were the case, the non-recombining sex-linked region already had a low recombination rate in the proto-SD region prior to the acquisition of the SD gene (He et al. 2021). The low recombination rate and gene density of the PCH region thus provides excellent conditions for the SD gene to arise, without the costly need to create a new non-recombining region in the genome from scratch. Without further expansion of the non-recombining region, the sex chromosome can remain homomorphic as it was in the proto-sex chromosomes in face of occasional recombination between sex chromosomes (Stöck et al. 2011; Rodrigues et al. 2018).

**Fig. 5.**
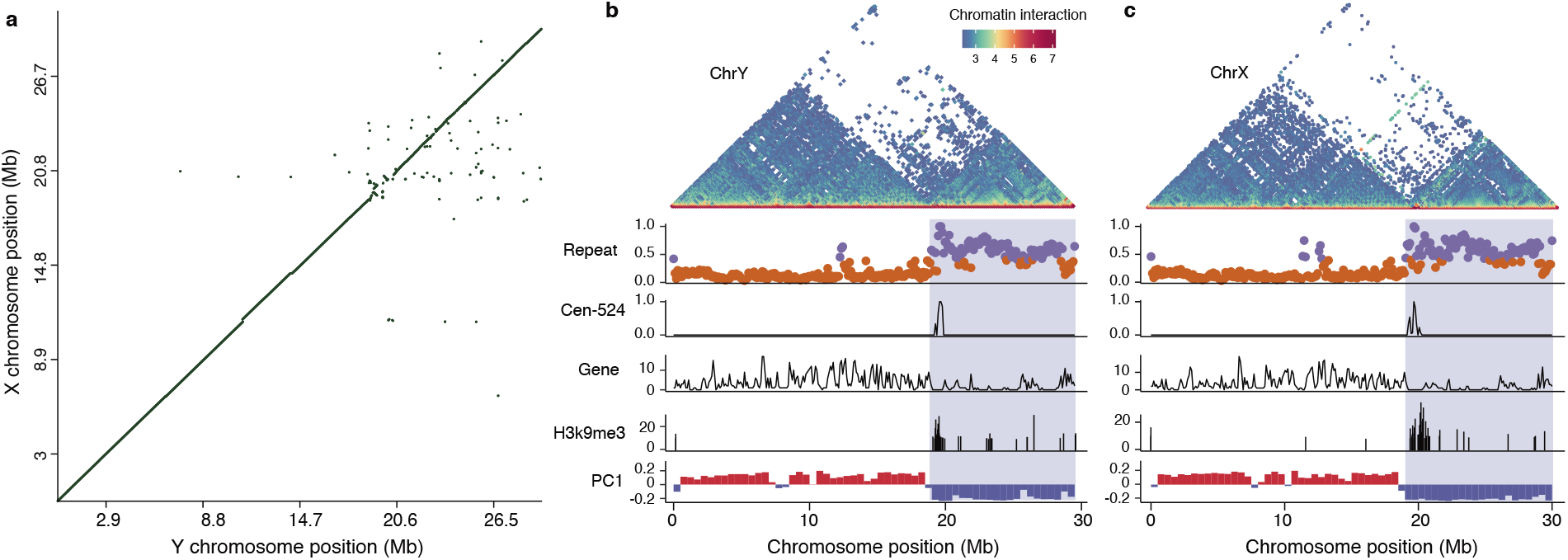
comparison of chromosome configuration between the sex chromosomes. **a)** sequence synteny between the X and Y chromosome. No large-scale inversions spanning the non-recombining region were observed. **b-c)** Similar pattern of chromosome compartmentalization between the X and Y chromosome. The legends for the Hi-C contact map, repeat content, distribution of Cen-524, gene density, H3K9me3 modification and A/B compartments are similar to these described in Fig. 3a-c. The PCH region is highlighted with blue background.

### Conclusions

The nearly complete diploid genome of zig-zag eels provides an opportunity to elucidate the sequences and organization of the centromeres and PCH (Lee et al. 2020). Despite the lack of direct imaging of subnuclear organization (Su et al. 2020; Takei et al. 2021; S. Wang et al. 2016), the Hi-C data uncover the pattern of compartmentalization that differ between PCH and euchromatin, allowing us to infer the possible spatial positioning of chromatin (J. Liu et al. 2021; Lieberman-Aiden et al. 2009; Lesne et al. 2014). It has been suggested that similar compartments tend to interact with each other and the centromeres in many taxa clusters form chromocenter (Simon et al. 2015). We propose the position of centromeres and PCH can drive the spatial organization of the entire chromosome, but the impact is dependent on the size and type of chromosomes. Submetacentric and metacentric chromosomes seem to be more extensively subject to the driving force of PCH on compartmentalization, in part because the PCH takes up a larger proportion in relative to telocentric. However, it seems in the telocentric chromosomes the PCH spread from the centromere to a similar range, leading to smaller chromosomes having a larger portion of PCH. As a consequence, small telocentric chromosomes are also frequently influenced by the PCH on compartmentalization. It is unclear, however, whether the PCH associated chromosomal arms are positioned near the lamina or surround the nucleoli (van Steensel and Belmont 2017). Identifying the lamina-associated domains (LADs) and nucleolus-associated domains (NADs) will provide a more complete picture of the spatial distribution of PCH in the cell nuclei. Moreover, how the PCH-driven compartmentalization impacts gene regulation in the zig-zag eel remains elusive. Epigenetic profiling from multiple tissues with various markers other than H3K9me3 should promote the understanding of the regulatory landscape and mechanisms of gene expression across the chromosome (Yang et al. 2020; Robson, Ringel, and Mundlos 2019).

It should be noted that the characterization of PCH and its role in genome compartmentalization is limited to zig-zag eel in this study. It’s unclear how pervasive the PCH can take up a large portion of the genome and drive the organization of compartments, partly because in very few organisms the full sequences of centromere and pericentromeres have been assembled. Nevertheless, in zig-zag eel we identified one of the evolutionary consequences of the large PCH, that is, creating an condition (low recombination) for a sex-linked region to arise. A similar scenario has been reported in blue tilapia in which the sex-linked region locate on LG3 is highly repetitive and heterochromatinized in both the Z and W chromosome (Tao et al. 2021; Conte et al. 2020). This is in contrast with previous understanding of the origin of heterochromatin on the sex chromosome (Bachtrog 2005; Charlesworth 2017). Recent studies on ratites, an bird group with unusual homomorphic sex chromosome, suggested that the change of chromatin configuration and recombination rate can occur prior to sequence differentiation of the sex chromosomes (J. Liu et al. 2021; Xu et al. 2019). Expanding the study to more taxa with young sex chromosomes with the complete genome decoded will provide more understandings of the role of chromatin organization in sex chromosome origin and evolution.

## Methods

### Sample collection and genome sequencing

Total DNA was extracted from muscle tissue of one male fish using a QIAamp DNA Blood Mini Kit (Qiagen) according to the manufacturer’s instructions. Large-insert single-molecule real-time circular consensus sequencing (HiFi CCS) library preparation was conducted following the Pacific Biosciences recommended protocols. In brief, a total of 50 μg genomic DNA was sheared to ~20 kb targeted size by using Covaris g-TUBEs (Covaris). The sheared genomic DNA was examined by Agilent 2100 Bioanalyzer DNA12000 Chip (Agilent Technologies) for size distribution. Sequencing libraries, were constructed using the PacBio DNA template preparation kit 2.0 (Pacific Biosciences of California, Inc., Menlo Park, CA) for HiFi sequencing on the PacBio RS II machine (Pacific Biosciences of California, Inc.) according to the manufacturer’s instructions. The constructed libraries were sequenced on one SMRT cell on a PacBio RSII sequencer.

### RNA-seq

The total RNA was isolated from eye, brain, skin, testis, ovary, liver, spleen, kidney, intestines, muscle, blood, fin, gill, heart and pituitary gland tissues using the EasyPure RNA Kit (Transgen). Sequencing libraries were generated using the NEBNext® UltraTM RNA Library Prep Kit for Illumina® (NEB, Ipswich, MA, USA), following the manufacturer’s recommendations. the cDNA libraries used for paired-end (2 × 125 bp) sequencing on an Illumina HiSeq Xten platform by Annoroad Gene Technology Co. Ltd. (http://www.annoroad.com).

### Haplotype-resolved genome assembly

We used the peregrine (0.1.6.1) (Chin and Khalak 2019) assembler to assemble the ccs reads, with default parameters. The assembled contigs were aligned to a chromosome-level assembly of zig-zag eel GCF_900324485.2 (Rhie et al. 2020), and a pseudo-chromosome assembly of the contigs was generated by the Ragoo (1.1) (Alonge et al. 2019) program. The CCS reads were then mapped to the pseudo-chromosome assembly, using minimap2 (2.15-r905) (Heng Li 2018) with the parameter ‘ -k 19 -O 5,56 -E 4,1 -B 5 -z 400,50 -r 2k --eqx --secondary=no’. Each of the Hi-C read pairs sequenced from the same individual was mapped to the genome with BWA-MEM (0.7.16a) using the default parameters. Then the alignments of both reads were paired using the script HiC_repair from the hapCUT2 (v1.2) (Edge, Bafna, and Bansal 2017) package. To partition the haplotype-specific reads, we used a pipeline described in (Garg et al. 2020). Briefly, the CCS alignments were phased with Whatshap (0.18) (Patterson et al. 2015) and hapCUP2, using the phasing information derived from the CCS reads themselves, as well as the Hi-C read pairs. The two phased haplotypes were named hap-X and hap-Y respectively. We then partitioned the reads mapping to the hap-X and hap-Y blocks. The hap-X and hap-Y derived reads were then used for haploid genome assembly. The unphased reads, despite only a small fraction of the total reads, were added to both the hap-X and hap-Y reads for genome assembly. The peregrine assembler was again used for haploid genome assembly

### Chromosome level assembly

We combined the two haploid genomes together as a diploid reference against which we mapped the Hi-C reads pairs. Each of the Hi-C read-pair was mapped to the reference using BWA-MEM with parameters “-A 1 -B 4 -E 50 -L 0”. We then extracted the read-pairs that were mapped to hap-X and hap-Y contigs respectively. To do so, we required each of the Hi-C read pairs to have zero mismatch, and both of the read pairs were mapped to either hap-X or hap-Y contigs. The read-pairs mapped to the hap-X and hap-Y were used to do scaffolding for hap-X and hap-Y contigs respectively. We used the 3D-DNA (180114) pipeline (Benson 1999; Dudchenko et al. 2017) to scaffold the contigs. The Hi-C reads were mapped to the haploid genome using the Juicer (1.7.6) pipeline (Durand, Shamim, et al. 2016). The alignment information was used by the 3D-DNA pipeline for scaffolding and producing the final Hi-C contact map. The Hi-C contact map was then visualized by Juicebox (1.11.08) (Durand, Robinson, et al. 2016) which allowed for manual curations. We corrected inversion errors in the Juicebox interactive interface and re-joined contigs that failed to be linked by 3D-DNA.

### Genome annotation

We used RepeatModeler to predict new repeat families in the zig-zag eel genome. To annotate tandem repeats, we searched for candidate repeat units using Tandem Repeat Finder (Benson 1999) with the parameter “2 7 7 80 10 20 2000 -l 6”. The results were filtered by pyTanFinder (Kirov et al. 2018) which removed the redundancy of the repeat units. The predicted new repeat families and tandem repeats were combined with the existing fish repeat library from Dfam (3.1) and RepBase (20170127) as the input library for repeat annotation and masking. We used the Liftoff (1.2.1) (Shumate and Salzberg 2020) program to translate the existing RefSeq gene model annotations (GCF_900324485.2) into the new assemblies of both haplotypes, with default parameters.

### Phylogenomics

We used Last (1042) (Kiełbasa et al. 2011) to align the genomes of seven species, swamp eel (Monopterus albus) (Zhao et al. 2018), Nile tilapia (*Oreochromis niloticus*) (Tao et al. 2021), yellow perch (*Perca flavescens*) (Feron et al. 2020), threespine stickleback (*Gasterosteus aculeatus*) (Peichel et al. 2020), yellowbelly pufferfish (*Takifugu flavidus*), big-bellied seahorse (*Hippocampus abdominalis*) and pricklefish (*Acanthochaenus luetkenii*) (Musilova et al. 2019), against the hap-Y reference genome with the option “-m100 -E0.05 -C2”. The one-to-one best alignments were retained to build 7-way multiple alignments using MULTIZ (v11.2) (Blanchette et al. 2004). We then reconstructed the phylogenetic tree using IQTREE (2.0-rc1) (Minh et al. 2020) with 1000 times bootstrapping. The inferred phylogeny was used for estimation of species divergence time with Beast (2.6.0). The range of 98.0 - 100.5 million year was set for age calibration according to the fossil records of Acanthopterygii (Musilova et al. 2019).

### FISH experiment

Amplification of the centromeric (Cen-524) and telomeric (Tel-190) sequences was conducted using the primers listed in the Supplementary Table S2. PCR amplification was performed in a 50-μL volume containing 1× ExTaq buffer, 500 nM primers, 200 μM dNTPs and 1.25 U ExTaq DNA polymerase (TaKaRa Bio, Kusatsu, Shiga, Japan). PCR was conducted using the following parameters: denaturation at 95 °C for 3 min; 35 amplification cycles of 95 °C for 30 s; 55 °C for 30 s; 72 °C for 30 s; and an extension at 72 °C for 10 min. The PCR products were checked using agarose gel electrophoresis, and then purified and eluted in water using a QIAquick PCR purification kit (Qiagen, Hilden, Germany). The purified PCR products of Cen-524 and Tel-190 were labeled with Cy5-dUTP and FITC-dUTP using Nick Translation Mix (Roche, Mannheim, Germany), respectively. Nick translation was performed at 15°C for 1.5 h. These two probes were checked via agarose gel electrophoresis.

Chromosomal preparation of *M. armatus* was used for a FISH experiment as previously described (J. D. Liu et al. 2002). Briefly, the slides were treated with 0.01% pepsin solution in 0.1 N HCl at 37°C for 1 h, and then a 5-min wash using 2× SSC was performed. Next, they were fixed for 1 min in 4% formaldehyde in 2× SSC, rinsed three times for 3 min in 2× SSC, dehydrated with 70%, 95%, and 100% ethanol series at room temperature for 3 min each, and finally air-dried. The chromosome preparations were denatured in 70% formamide for 1 min at 70°C. The slides were dehydrated in 70%, 95% and 100% ethanol series at −20°C, 5 min each. The 20 μl of the hybridization mix contains 50% formamide, 10 mg/mL dextran sulfate, 2× SSC and 100 ng of each probe was heated to 95°C for 10 min and stored at 4 °C until use. Hybridization was performed during 16 h at 37°C. After hybridization, slides were washed three times for 5 min in 2× SSC, then washed again in 1 × PBS at room temperature for 5 min. Slides were mounted with DAPI (Vector Laboratories, Odessa, Florida, United States). Chromosomes and FISH signals were visualized with an Olympus BX63 fluorescence microscope (Olympus, Tokyo, Japan). Images were captured on cellSens Dimension v. 1.9 and an Olympus DP80 CCD camera (Olympus, Tokyo, Japan). Images were adjusted with Adobe Photoshop v. 8.0 (Adobe, San Jose, CA, USA).

### Hi-C analysis

We mapped the haploid-specific Hi-C read pairs to the corresponding haploid assemblies using the Juicer pipeline mentioned above. To generate the Hi-C contact matrix, we applied the ‘pre’ function of juicebox_tools from the Juicer package. Only intra-chromosomal interactions were calculated. Then we extracted the count values by applying the ‘dump’ function for each chromosome, normalizing the data with the KR method. The interaction frequency was calculated with a bin size of 50 kb. For each pair-wise interaction, we required the count number larger than 10. When visualizing the chromosome-wide matrix, we further used the log transformed matrix. To call the compartments, we applied the ‘eigenvector’ function of juicebox_tools with KR normalization. We calculated the eigenvector at 250 kb resolution. The first principal component of the Pearson’s correlation matrix based on the intrachromosomal matrix was used as the eigenvectors.

### H3K9me3 histone modification

CUT&Tag assay was performed as described previously with modifications (Kaya-Okur et al. 2019). Briefly, native nuclei were purified from frozen samples as previously described (Corces et al. 2017) and were washed twice gently with wash buffer (20 mM HEPES pH 7.5; 150 mM NaCl; 0.5 mM Spermidine; 1 × Protease inhibitor cocktail). A 1:50 dilution of H3K9me3 (ab8898) or IgG control antibody (normal rabbit IgG: Millipore cat. no. 12-370) was used for incubation. The secondary antibody (Anti-Rabbit IgG antibody, Goat monoclonal: Millipore AP132) was diluted to 1:100 in the dig wash buffer. DNA was purified using the phenol-chloroform-isoamyl alcohol extraction and ethanol precipitation. The libraries were amplified by mixing the DNA with 2μL of a universal i5 and uniquely barcoded i7 primer. The size distribution of libraries was determined by Agilent 4200 TapeStation analysis. Sequencing was performed in the Illumina Novaseq 6000 using 150bp paired-end following the manufacturer’s instructions. We followed the bioinformatic pipeline described in (Kaya-Okur et al. 2019) to process the sequencing reads. Briefly, we used Bowtie2 (2.3.5.1) (Langmead and Salzberg 2012) to map the raw reads against the genome with the options “--local --very-sensitive-local --no-unal --no-mixed --no-discordant -I 10 -X 700”. Read redundancy was removed by the rmdup command of SAMtools (1.11) (H. Li et al. 2009). We used macs2 (2.2.7.1) (Zhang et al. 2008) to call peaks, and selected peaks with - log10 p-values larger than 8.

### Sex-linked regions

We collected re-sequencing data from ten males and ten females. The sex of zig-zag eel was identified through histological analysis of gonad. Once the sex was identified, we extract DNA from the individual muscle tissue. The extracted DNA molecules were sequenced with Illumina X Ten platform (San Diego, CA, USA), a paired-end library was constructed with an insert size of 250 base pairs (bp) according to the protocol provided by the manufacturer. The raw reads were mapped against the genome with BWA-MEN using the default parameters. After marking the duplicates, we call single-nucleotide polymorphic sites (SNPs) with the GATK (4.1.4.0) joint calling pipeline. To filter the variants, we applied the parameters “QD < 2.0 || FS > 60.0 || MQRankSum < −12.5 || RedPosRankSum < −8.0 || SOR > 3.0 || MQ < 40.0”. We used SNPeff to select the SNPs that are heterozygous in one sex but homozygous in the other sex. We further used the EMMAX (8.22) pipeline to identify the SNPs that are associated with sex.

### Gene expression

The RNA-seq data of gonads from five males, five females and five intersex individuals was produced in Xue et al. (submitted). The raw reads were mapped against the genome using HiSat2 (2.1.0) with the option “-k 4”. The numbers of reads that map to transcripts of coding genes were counted with featureCount using default parameters. The expression levels were quantified using the TPM (transcripts per million) value.

### Recombination rate

We used the ReLERNN (1.0.0) method to estimate recombination rates using the resequencing data of 20 individuals. We used the filtered variants mentioned above with the non-biallelic variants further removed. Since the sex chromosomes are not diploid, they are excluded from analyses. We simulated and trained the datasets with the default parameters. Finally, we estimated the recombination rates in non-overlapping windows whose sizes were decided by ReLERNN.

## Data availability

The genome assemblies and sequencing data are deposited at NCBI under the accession PRJNA608290. A full list of accession IDs is available in the Supplementary Table S3.

## Code availability

The scripts used in this study have been reposed at Github (https://github.com/lurebgi/zigzigEelSexchr/).

## Acknowledgement

This work was supported by the Special Fund for Agro-Scientific Research in the Public Interest of Fujian Province of China (2020R1014002) and the Fujian Provincial Marine and Fishery Structure Adjustment Fund (2020MDS-YT005) to LZX, the National Natural Science Foundation of China (32002360) and the Natural Science Foundation of Yunnan Province of China (2018FB048) to YG. LHX. is supported by the Erwin Schrödinger Fellowship (J4477-B) from the Austrian Science Fund (FWF).

