## Supplementary figure for "Pericentromeric heterochromatin impacts genome compartmentalization and sex chromosome evolution in a fish": Supplementary figure.docx

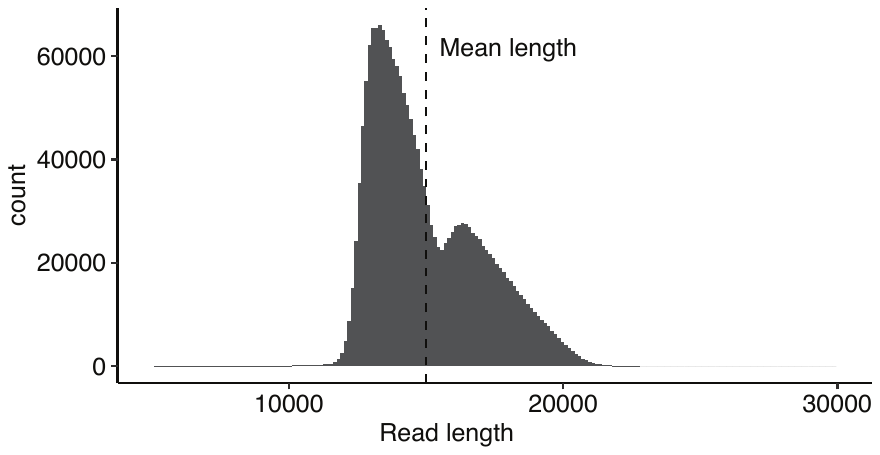

**Supplementary Fig. S1 The distribution of HiFi read lengths**.

The length of all HiFi reads were calculated and their distribution was shown. The vertical dashed line shows the mean length of CCS reads.

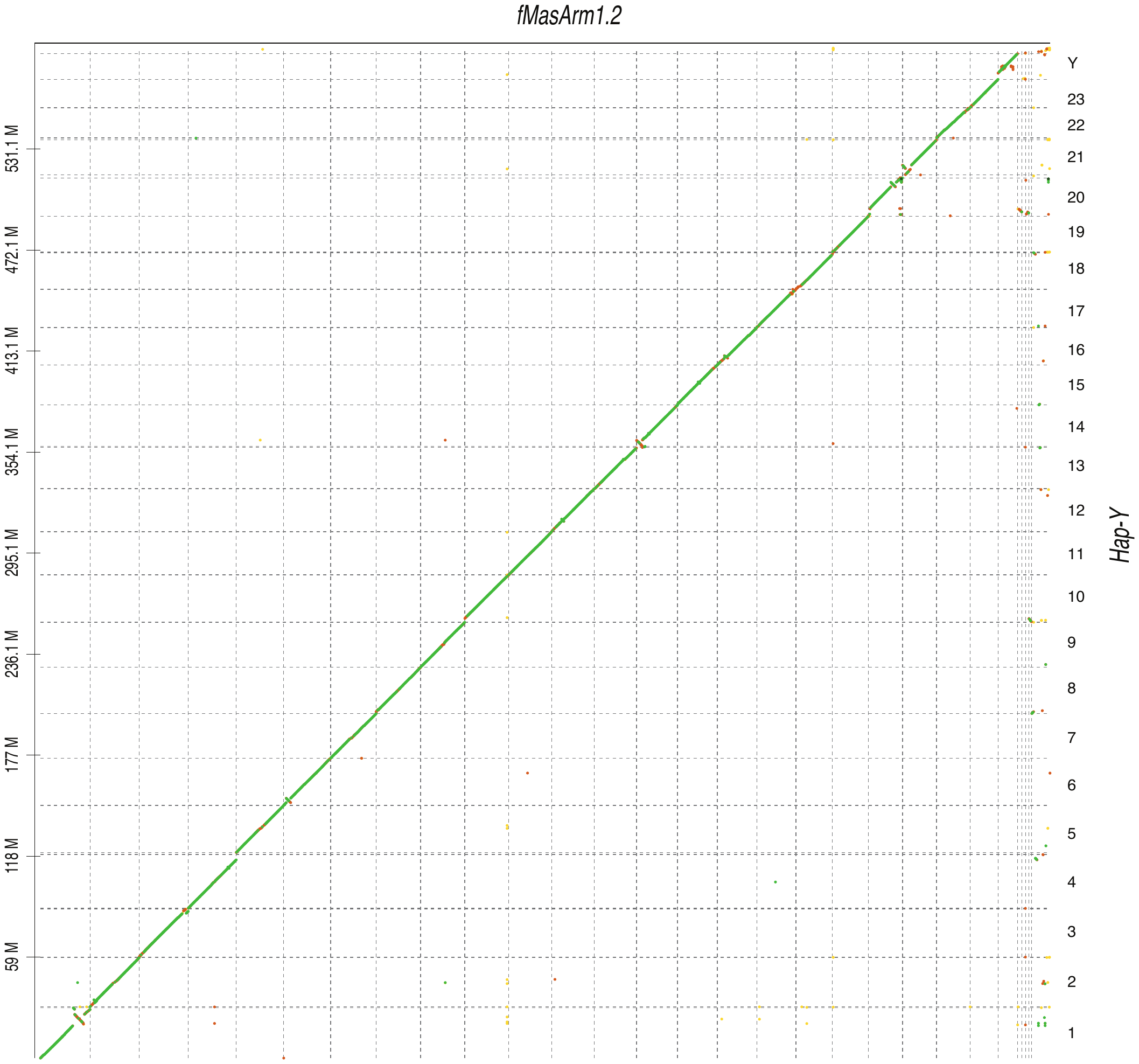

**Supplementary Fig. S2 The dot-plot between the assembly of Hap-Y and fMasArm1.2**. Hap-Y is the haploid genome of zig-zag eel produced in this study the fMasArm1.2 is the genome assembly produced by Vertebrate Genome Project (VGP). The reddish colors indicate low sequence similarity while the green colors indicate high sequence similarity.

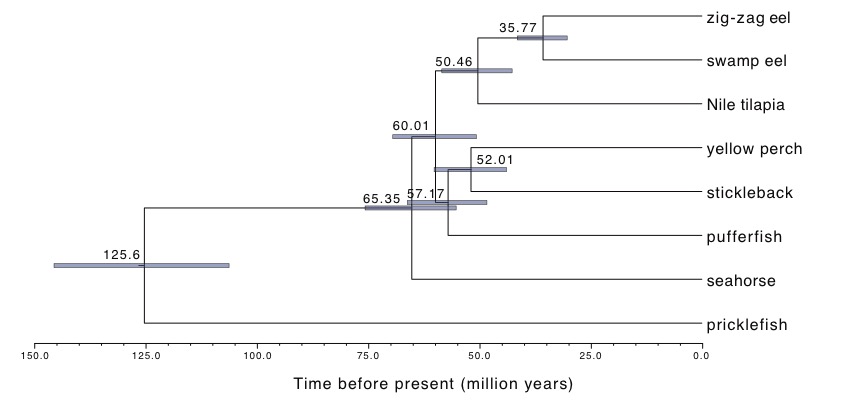

**Supplementary Fig. S3 Dating of species divergence in Percomorpha**.

Whole genome alignments were used to estimate species divergence. The estimated ages were calibrated with the fossil record at the ancestor node (Acanthopterygii). The error range shows the 95% confidence interval.

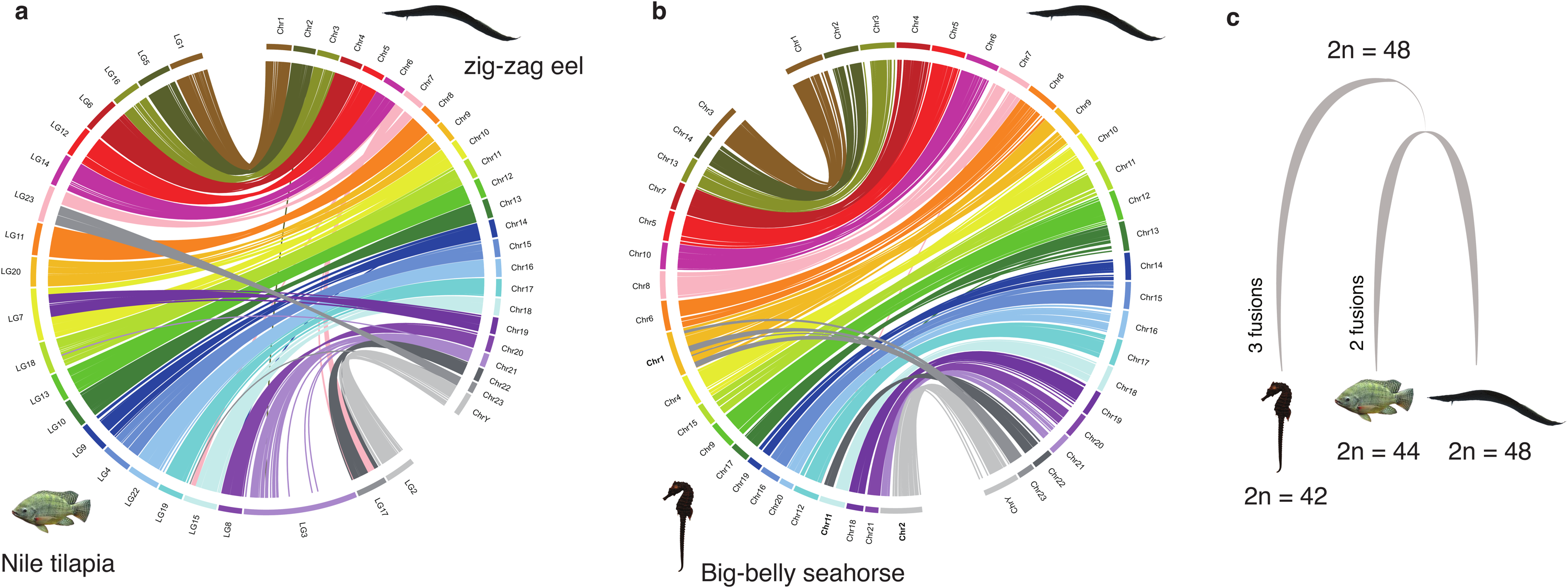

**Supplementary Fig. S4 Reconstruction of Percomorpha ancestral karyotype**.

**a-b**) The chromosome synteny between the zig-zag eel and two other fishes: Nile tilapia and big-belly seahorse. **c**) A schematic diagram shows the changes of chromosome number during the evolution of Percomorpha species. The diploid number (2n) was shown for each species. The occurrence and times of fusions were inferred based on the parsimonious principle.

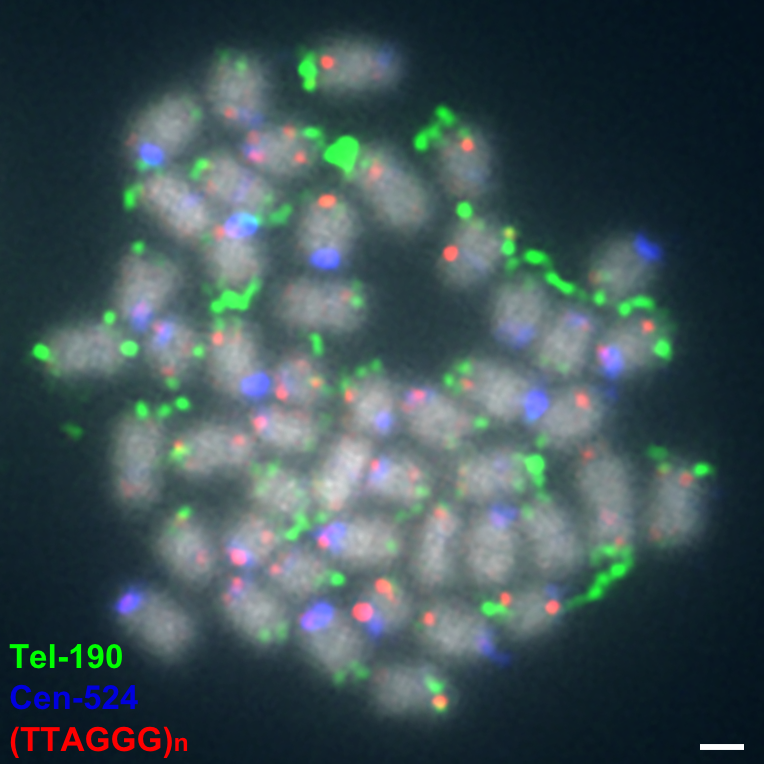

**Supplementary Fig. S5 FISH mapping of Cen-524 and Tel-190 probes on somatic metaphase chromosomes**.

Clear signals of Cen-524 were detected in most of the centromeric regions, and clear signals of Tel-190 were detected in the telomeric regions of one chromosome arm on most somatic metaphase chromosomes. In most Chromosomes the conserved telomere motif (TTAGGG)n are present and sometimes are co-present with Tel-190. Scale bars = 1 μm.

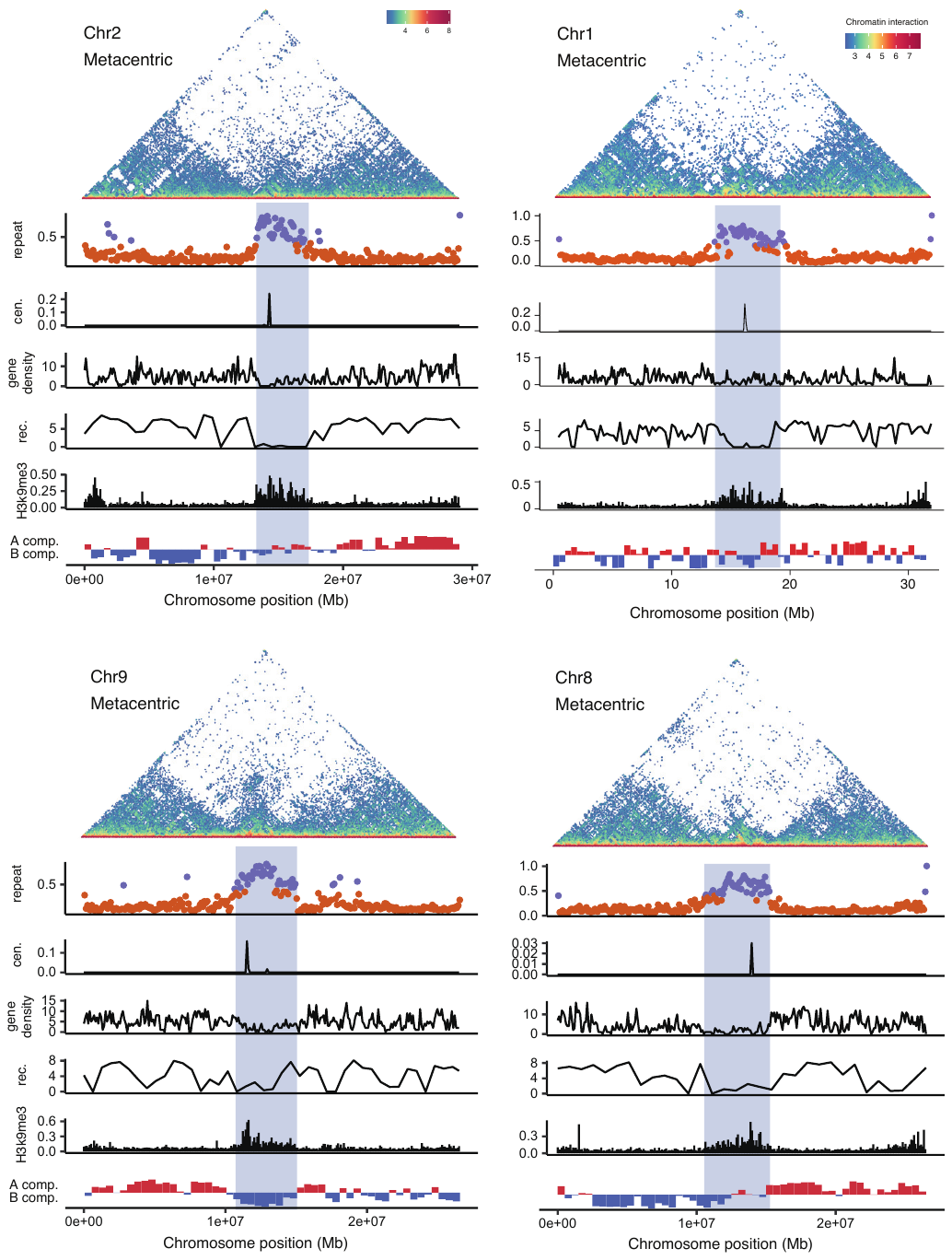

**Supplementary Fig. S6. Identification of PCH on metacentric chromosomes.**

In the top panel, the colors of dots measure the frequency of chromatin interacting between 100 kb windows. When the repeat content of a 50 kb sequence (a dot) is larger than 40%, it is highlighted in dark purple, otherwise in orange. The portion (%) of Cen-524 satellite in 100 kb windows. The gene density is measured as the number of genes in 100 kb windows. The recombination rate (Rec.) is estimated with selected window size based on the available variants. The Y-axis of the H3K9me3 panel shows the -log 10 transformed p-values for the H3K9me3 peaks. The PC1 panel shows the PC1 values of Hi-C epivector: the positive values (red) represent active (A) compartments and the negative values (blue) represent silenced (B) compartments.

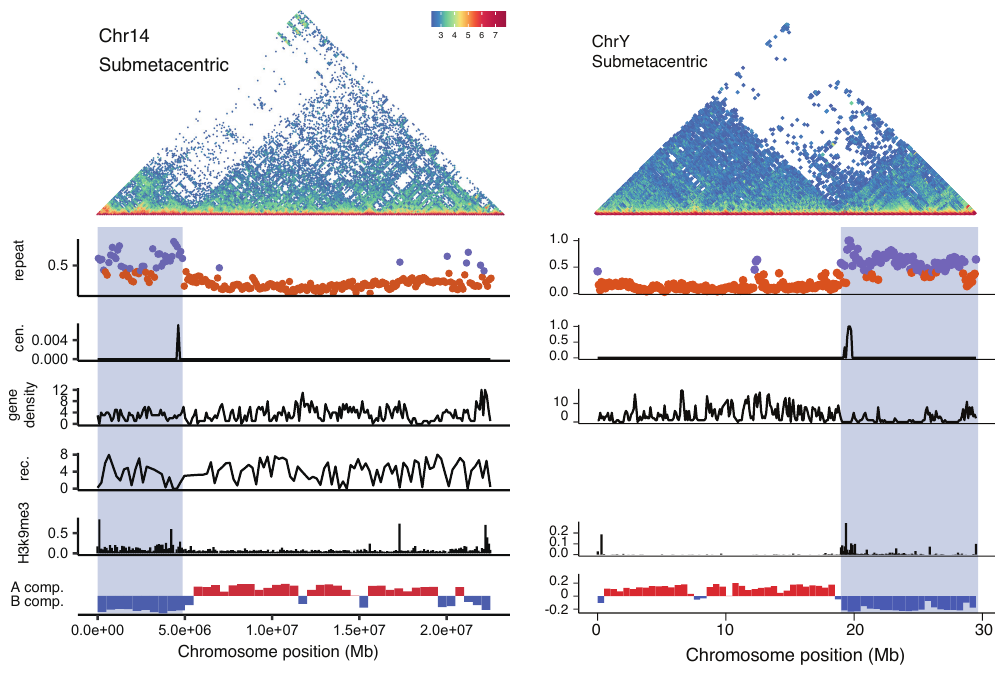

**Supplementary Fig. S7. Identification of PCH on submetacentric chromosomes.**

We were unable to estimate the recombination rate for the Y chromosomes using the population data. In the top panel, the colors of dots measure the frequency of chromatin interacting between 100 kb windows. When the repeat content of a 50 kb sequence (a dot) is larger than 40%, it is highlighted in dark purple, otherwise in orange. The portion (%) of Cen-524 satellite in 100 kb windows. The gene density is measured as the number of genes in 100 kb windows. The recombination rate (Rec.) is estimated with selected window size based on the available variants. The Y-axis of the H3K9me3 panel shows the -log 10 transformed p-values for the H3K9me3 peaks. The PC1 panel shows the PC1 values of Hi-C epivector: the positive values (red) represent active (A) compartments and the negative values (blue) represent silenced (B) compartments.

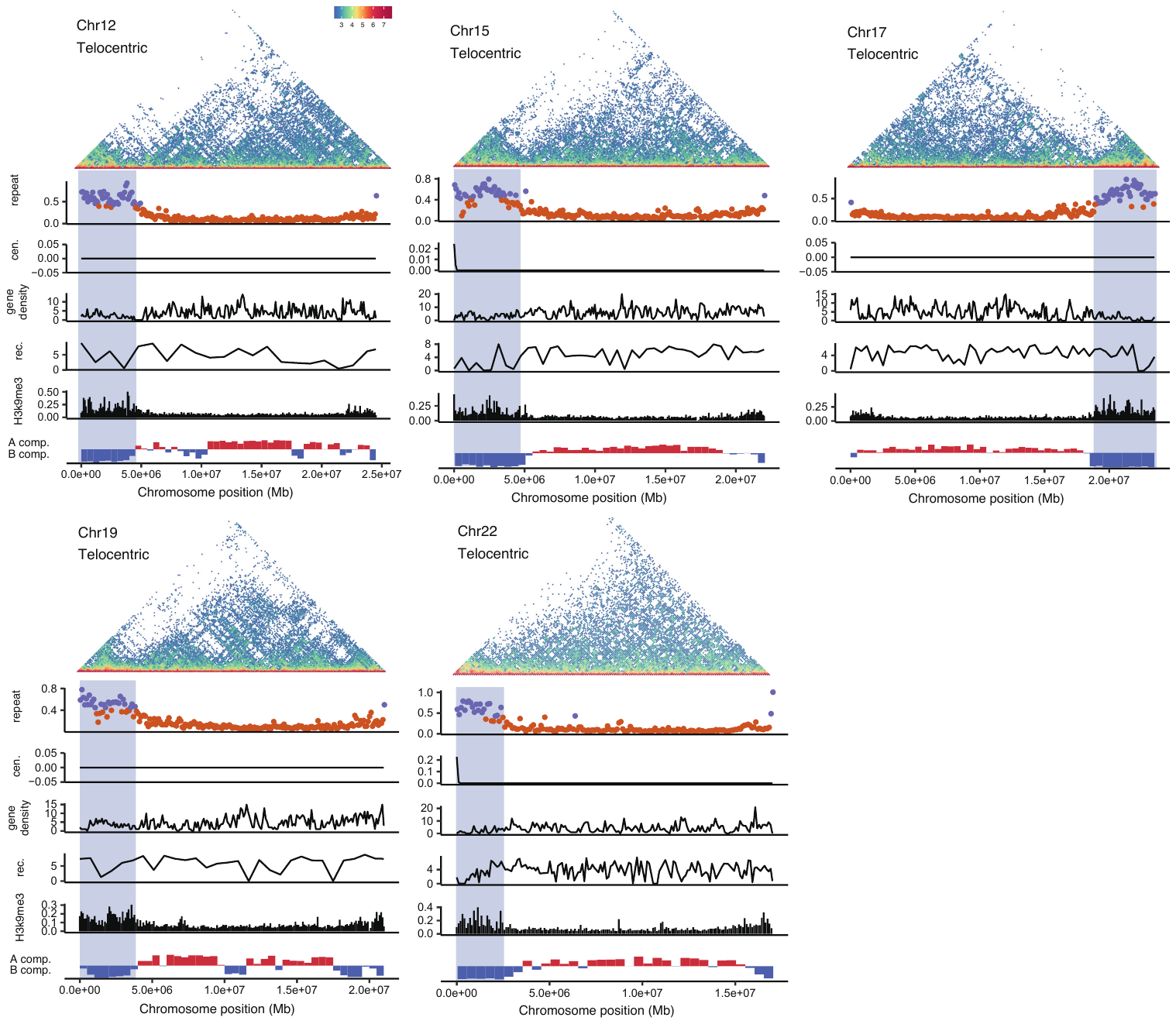

**Supplementary Fig. S8. Identification of PCH on small telocentric chromosomes.**

In the top panel, the colors of dots measure the frequency of chromatin interacting between 100 kb windows. When the repeat content of a 50 kb sequence (a dot) is larger than 40%, it is highlighted in dark purple, otherwise in orange. The portion (%) of Cen-524 satellite in 100 kb windows. The gene density is measured as the number of genes in 100 kb windows. The recombination rate (Rec.) is estimated with selected window size based on the available variants. The Y-axis of the H3K9me3 panel shows the -log 10 transformed p-values for the H3K9me3 peaks. The PC1 panel shows the PC1 values of Hi-C epivector: the positive values (red) represent active (A) compartments and the negative values (blue) represent silenced (B) compartments.

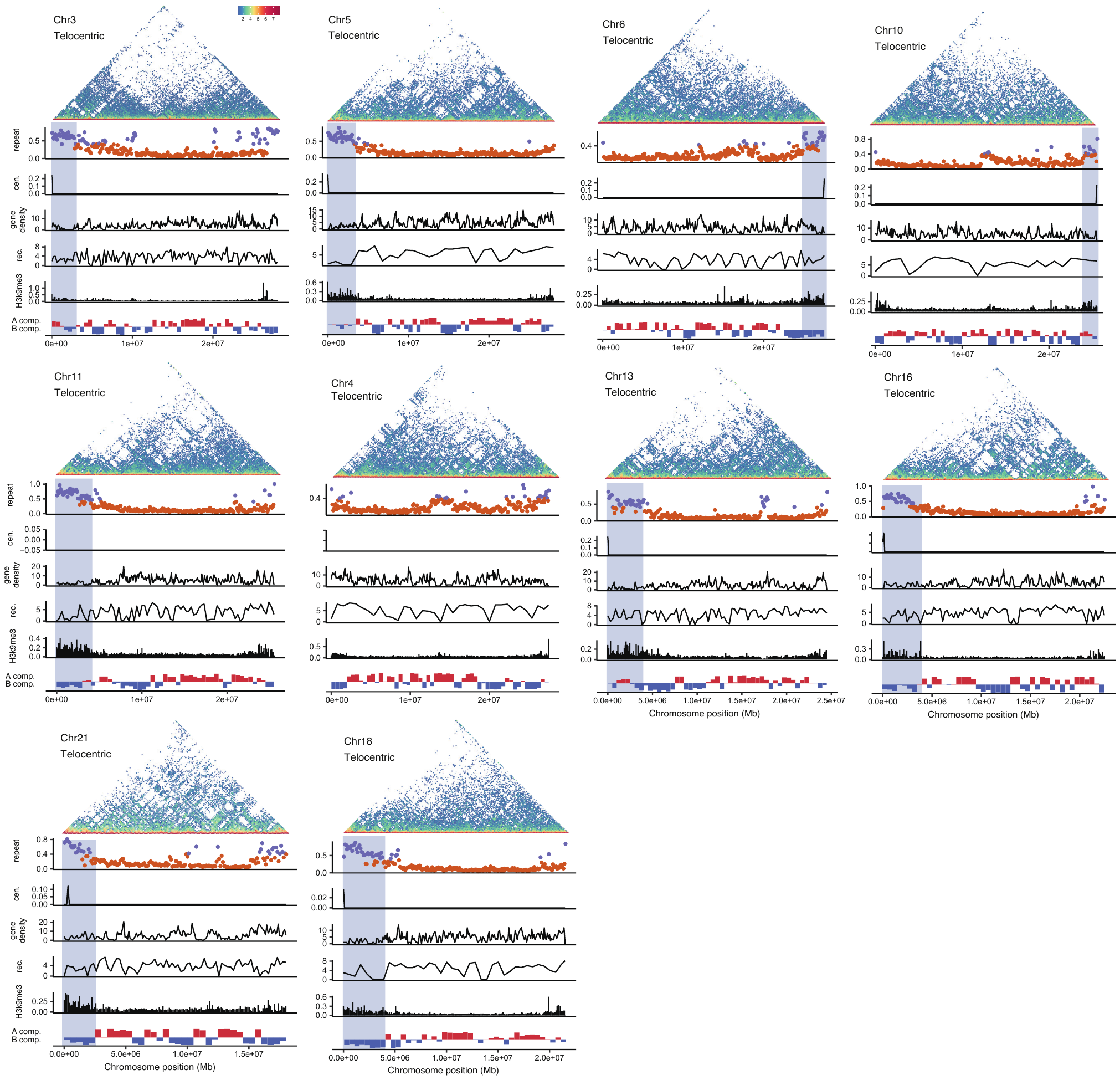

**Supplementary Fig. S9. Identification of PCH on large telocentric chromosomes.**

In the top panel, the colors of dots measure the frequency of chromatin interacting between 100 kb windows. When the repeat content of a 50 kb sequence (a dot) is larger than 40%, it is highlighted in dark purple, otherwise in orange. The portion (%) of Cen-524 satellite in 100 kb windows. The gene density is measured as the number of genes in 100 kb windows. The recombination rate (Rec.) is estimated with selected window size based on the available variants. The Y-axis of the H3K9me3 panel shows the -log 10 transformed p-values for the H3K9me3 peaks. The PC1 panel shows the PC1 values of Hi-C epivector: the positive values (red) represent active (A) compartments and the negative values (blue) represent silenced (B) compartments.

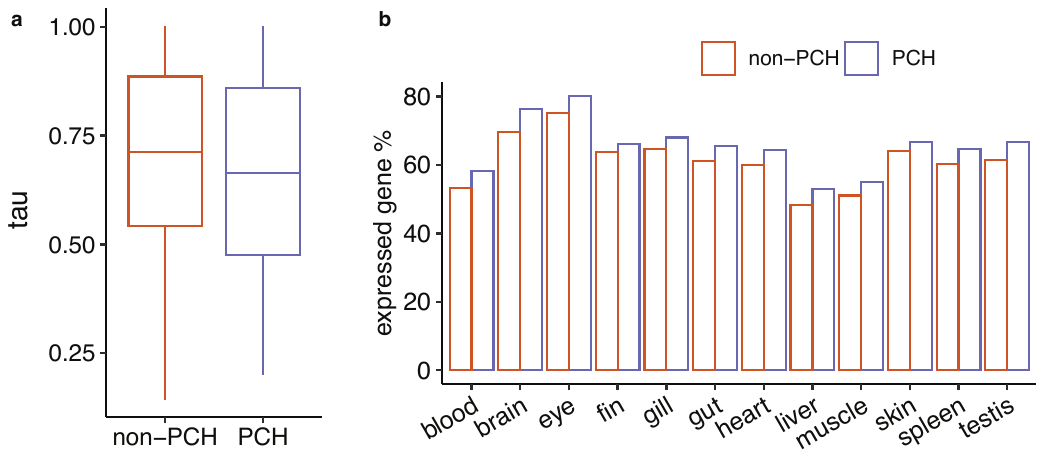

**Supplementary Fig. S10 PCH contains more active genes which are expressed more broadly.**

**a)** The tau index for genes in PCH and non-PCH regions. PCH genes are more broadly expressed than non-PCH genes (P = 2.279e-11, Wilcoxon rank sum test). A lower value of tau means larger breadth of expression. **b)** A larger proportion of expressed genes in PCH than in non-PCH regions. The expressed genes were defined as those with TPM (transcript per million) larger than 1.

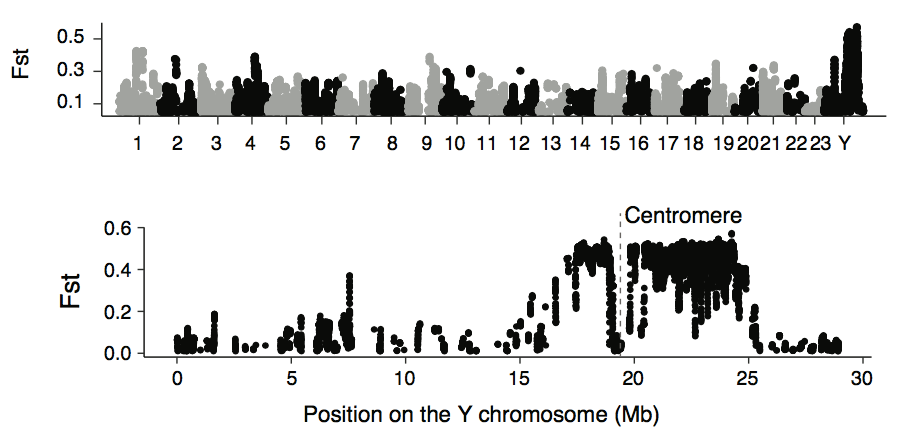

**Supplementary Fig. S11 population differentiation between the sexes is largest in the sex-linked region.**

The index of population differentiation (Fst) was calculated in 50 kb windows. The peaks of Fst are enriched on the Y chromosome. In the lower panel, the zoom-in view for the Y chromosome is shown. In the sex-linked region (17-24 Mb), the Fst values are the largest. We filtered out the windows that contain less than 40 variants.
